# Engineering TALE-linked deaminases to facilitate precision adenine base editing in mitochondrial DNA

**DOI:** 10.1101/2023.09.03.556141

**Authors:** Sung-Ik Cho, Kayeong Lim, Seongho Hong, Jaesuk Lee, Annie Kim, Ji Min Lee, Young Geun Mok, Eugene Chung, Seunghun Han, Sang-Mi Cho, Jieun Kim, Sanghun Kim, Eun-Kyoung Kim, Ki-Hoan Nam, Yeji Oh, Minkyung Choi, Seonghyun Lee, Hyunji Lee, Jin-Soo Kim

**Affiliations:** Center for Genome Engineering, Institute for Basic Science, Daejeon, 34126, Republic of Korea; Department of Chemistry, Seoul National University, Seoul, 08826, Republic of Korea; NUS Synthetic Biology for Clinical & Technological Innovation (SynCTI) and Department of Biochemistry, National University of Singapore, Singapore; Laboratory Animal Resource and Research Center, Korea Research Institute of Bioscience and Biotechnology, Cheongju, 28116, Republic of Korea; Department of Bioscience, Korea University of Science and Technology, Daejeon, 34113, Republic of Korea; Department of Biochemistry, College of Natural Sciences, Chungbuk National University, Cheongju, 28644, Republic of Korea; College of Veterinary Medicine and Research Institute of Veterinary Medicine, Chungbuk National University, Cheongju, 28644, Republic of Korea; School of Medicine, Sungkyunkwan University, Suwon, 16419, Republic of Korea; Brain Science Institute, Korea Institute of Science and Technology, Seoul, 02792, Republic of Korea; Division of Bio-Medical Science & Technology, KIST School, Korea University of Science and Technology, Seoul, 02792, Republic of Korea; Edgene, Inc., Seoul, 08790, Republic of Korea; GreenGene Inc., Seoul, 08790, Republic of Korea; Department of Pharmacology, Yonsei University College of Medicine, Seoul, 03722, Republic of Korea

## Abstract

*DddA*-derived cytosine base editors (DdCBEs) and transcription activator-like effector (TALE)-linked deaminases (TALEDs) catalyze targeted base editing of mitochondrial DNA (mtDNA) in eukaryotic cells, a method useful for modeling of mitochondrial genetic disorders and developing novel therapeutic modalities. Here, we report that A-to-G editing TALEDs but not C-to-T editing DdCBEs induce tens of thousands of transcriptome-wide off-target edits in human cells. To avoid these unwanted RNA edits, we engineered the substrate-binding site in TadA8e, the deoxy-adenine deaminase in TALEDs, and created TALED variants with fine-tuned deaminase activity. Our engineered TALED variants not only reduced RNA off-target edits by > 99% but also minimized off-target mtDNA mutations and bystander edits at a target site. Unlike wild-type versions, our TALED variants were not cytotoxic and did not cause developmental arrest of mouse embryos. As a result, we obtained mice with pathogenic mtDNA mutations, associated with Leigh disease, which showed reduced heart rates.

## Main

Targeted base editing of mammalian mitochondrial DNA (mtDNA) is a powerful and versatile technology^1–7^, which can be used to model mitochondrial genetic diseases in cell lines and animals^8–15^ and to develop novel therapeutic modalities that could be used to correct pathogenic mutations in the future^16,17^. Programmable deaminases, which are composed of a custom DNA-binding protein and a nucleobase deaminase, enable mitochondrial DNA editing in a targeted manner^13, 18–25^. There are two types of programable deaminases that have been developed for organellar genome editing: cytosine base editors^13, 18, 20–24, 26^, including DddA-derived cytosine base editors (DdCBEs) and zinc finger deaminases, both of which induce C-to-T (= G-to-A on the other DNA strand) editing, and adenine base editors^19^, termed TALEDs, which catalyze A-to-G (= T-to-C) editing. These base editors bind to a target DNA site in the mitochondrial genome and deaminate cytosine or adenine to yield uracil or hypoxanthine, respectively, which is base-paired with adenine or cytosine, respectively, during DNA replication or repair. As a result, cytosine or adenine base editors induce C-to-T or A-to-G conversions, respectively, in a targeted manner. Because most mitochondrial genetic disorders are caused by transition mutations (purine-to-purine or pyrimidine-to-pyrimidine) rather than transversion mutations (purine-to-pyrimidine or vice versa), cytosine and adenine base editors, together, can correct ∼85% of pathogenic mtDNA mutations, including those associated with Leber Hereditary Optic Neuropathy (LHON), Mitochondrial Encephalomyopathy, Lactic Acidosis, Stroke-like episodes (MELAS), Leigh syndrome, etc (www.mitomap.org).

Unlike cytosine base editors, TALEDs contain TadA8e, a deoxy-adenine deaminase engineered from the *E. coli*-derived tRNA-specific TadA protein. TadA8e^27^ and related TadA variants^28, 29^ are also essential components in CRISPR RNA-guided adenine base editors (ABEs)^30^, widely used for A-to-G base editing of nuclear DNA. However, RNA-guided ABEs cannot be used for organellar DNA editing due to the challenge of delivering guide RNA into organelles^31^. In addition, TadA8e in ABEs has residual deaminase activity for RNA substrates, leading to unwanted, transcriptome-wide off-target base editing^32–38^. Here, we show that mtDNA-targeted TALEDs with a Mitochondrial Target Sequence (MTS) can also exhibit off-target RNA editing, leading to cytotoxicity and growth arrest in mouse embryos. We also present engineered TALEDs with novel mutations that minimize unwanted RNA editing, with which we were able to obtain mtDNA-edited mice with pathogenic mutations.

## RESULTS

### Mitochondrial DNA-targeted TALEDs induce transcriptome-wide off-target edits

We first investigated whether the DddA-split TALEDs (sTALEDs), in which each of the split DddA halves is fused to a TALE protein, targeted to the *COX3.1* or *ND1* site in the mitochondrial genome could cause unwanted, off-target RNA editing in human embryonic kidney 293T (HEK 293T) cells. To this end, we performed transcriptome-wide sequencing of total RNA isolated from cells at day 2 post-transfection. Whole transcriptome sequencing of two independent biological replicates showed that the two sTALEDs, composed of the left- or right-side TALE fused to the N-terminal DddA_tox_ half split at G1397 (L-1397N or R-1397N, respectively) and the right- or left-side TALE fused to the C-terminal DddA_tox_ half split at G1397 and TadA8e (R-1397C-AD or L-1397C-AD), induced base conversions at > 50,000 sites with a frequency of at least 7% (Fig. 1). The vast majority (> 99.8%) of these base conversions were A-to-G edits (in cDNA reverse transcribed from RNA) rather than C-to-T edits, indicating that an adenine deaminase rather than a cytosine deaminase was responsible for these single-nucleotide alterations. In contrast, DdCBEs targeted to the same sites or a single sTALED subunit (L-1397N or R-1397N), used as a negative control, did not induce these off-target RNA edits. The other subunit containing TadA8e (L-1397C-AD or R-1397C-AD) caused A-to-G conversions, which resulted from A-to-I alterations in RNA (Supplementary Fig. S1). We also found that monomeric TALEDs (mTALEDs) and dimeric TALEDs (dTALEDs), although they are, in general, less efficient for DNA editing than are sTALEDs, also caused RNA off-target edits (Supplementary Fig. S2).

**Figure 1.**
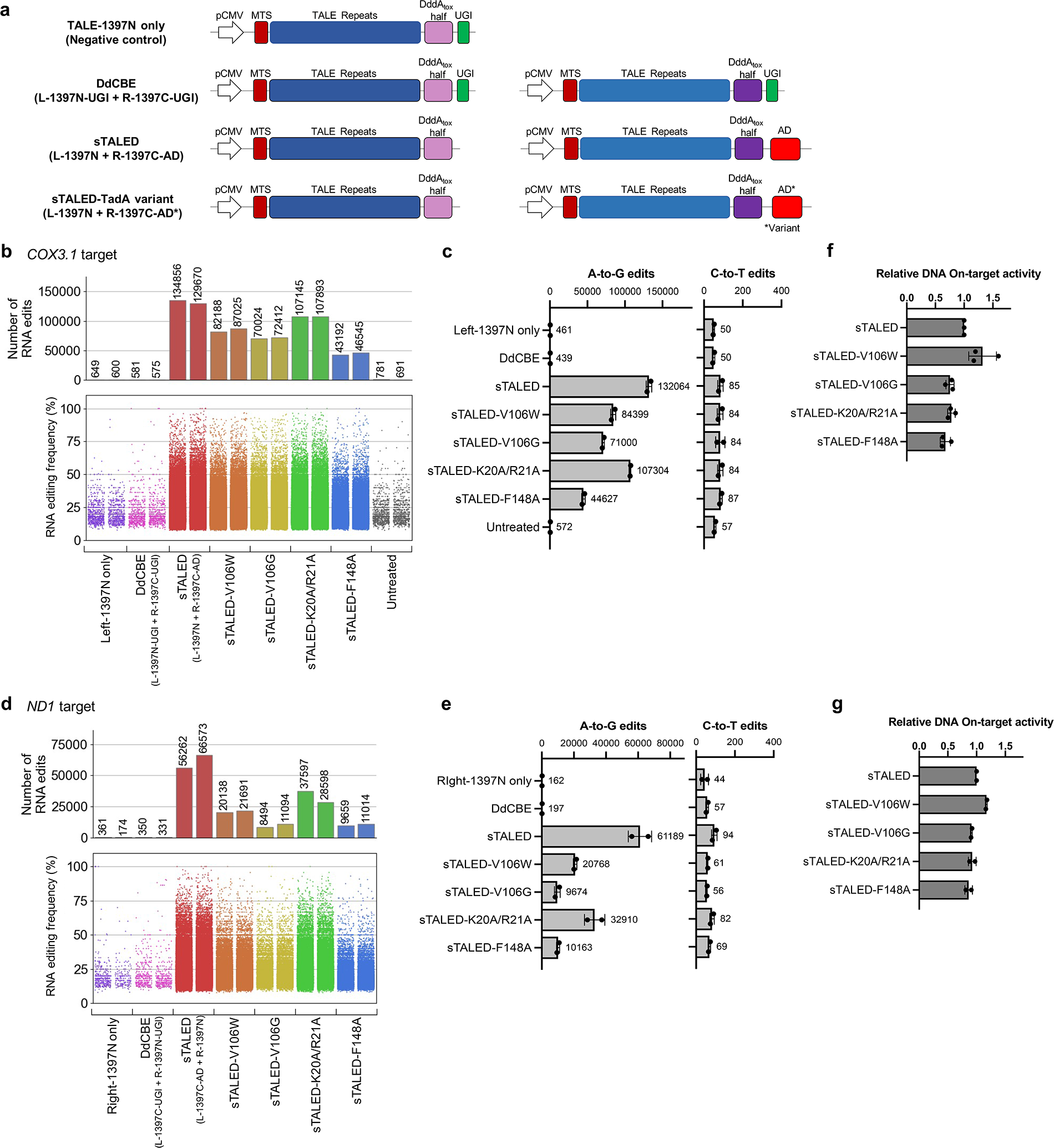
Transcriptome-wide A-to-I off-target edits induced by TALEDs. (a) Architecture of base editors used in the experiments. AD, TadA8e adenine deaminase; AD*, TadA8e adenine deaminase variant; MTS, mitochondrial targeting sequence; UGI, uracil glycosylase inhibitor. Light purple indicates the N-terminal halves of DddA_tox_ and dark purple indicates the C-terminal halves of DddA_tox_. (b-e) Bar graphs illustrating the total number of RNA edits found in HEK 293T cells that expressed DdCBE, sTALED, or sTALED variants containing previously described TadA variants developed for reducing RNA off-target editing (sTALED-V106W, sTALED-V106G, sTALED-K20A/R21A, and sTALED-F148A) as determined by whole transcriptome sequencing. Results from a negative control (cells expressing 1397N only) and untreated cells are also shown. (b) Number of RNA edits (top panel) and editing frequencies of each RNA edit for the *Cox3.1*-specific sTALEDs shown as a dot (bottom panel). (c) A-to-G edits and C-to-T edits for the *Cox3.1*-specific sTALEDs. (d) Number of RNA edits (top panel) and editing frequencies of each RNA edit for the *ND1*-specific sTALEDs shown as a dot (bottom panel). (e) A-to-G edits and C-to-T edits for the *ND1*-specific sTALEDs. Data are shown as means from n = 2 biologically independent samples. (f,g) The relative DNA on-target activity, normalized to that of the original sTALED, of the *Cox3.1*-specific sTALEDs (f) and *ND1*-specific sTALEDs (g). Data are shown as means from n=2 or 3 biologically independent samples. Source data are provided as a Source Data file.

To avoid or minimize unwanted, transcriptome-wide off-target A-to-I conversions induced by TALEDs, we introduced site-specific mutations in TadA8e, including V106W, V106G, K20A/R21A (dual mutations), or F148A, which are known to reduce off-target RNA editing^33–35^ when incorporated into CRISPR RNA-guided ABEs. Transcriptome-wide sequencing showed that sTALED variants incorporating these mutations in TadA8e reduced the number of off-target A-to-G edits significantly but not completely, while retaining DNA on-target editing efficiencies (Fig. 1). Thus, *Cox3.1*-specific sTALED variants with these site-specific mutations induced RNA off-target edits at sites that numbered from 44,627 (F148A) to 107,304 (K20A/R21A), reducing the number by 18.7% (= (132,064 – 107,304)/132,064) to 66.2% (= (132,064 – 44,627)/132,064), compared to the original sTALED (132,064 A-to-G edits) (Fig. 1b,1c and Supplementary Fig. S3). Likewise, *ND1*-specific sTALED variants reduced the number of RNA off-target edits by at least 46.2% (= (61,189 – 32,910)/61,189) (K20A/R21A) and up to 84.2% (= (61,189 – 9,674)/61,189) (V106G) (Fig. 1d,1e and Supplementary Fig. S4). These results show that the same TadA8e variants, used for reducing the RNA off-target editing effects of CRISPR RNA-guided ABEs, can also reduce collateral damage caused by sTALEDs. Yet, at least 9,600 and up to 107,000 off-target A-to-G edits persisted even with the best performing TadA8e variants.

### Protein engineering of TALEDs to avoid RNA off-target edits

To further minimize transcriptome-wide off-target edits, we sought to engineer TALEDs by mutating amino-acid residues, including V106, at the substrate binding site in TadA8e. To this end, we first chose 11 amino-acid residues at the substrate binding site, based on the 3D cryo-electron microscopy structure of ABE8e bound to DNA^39^ (Fig. 2a), and replaced each of these residues with the other 19 amino-acid residues in the *Cox3.1*-specific sTALED, making a total of 209 (= 11 X 19) sTALED variants. Plasmids encoding each of the resulting sTALED pairs were transfected into HEK 293T cells. We then measured on-target editing frequencies at the *Cox3.1* site at day 4 post-transfection. Among the 209 sTALED variants, a total of 101 sTALED pairs remained highly active in targeted mtDNA editing with an efficiency of at least 50%, compared to the wild-type sTALED pair (Supplementary Fig. S5).

**Figure 2.**
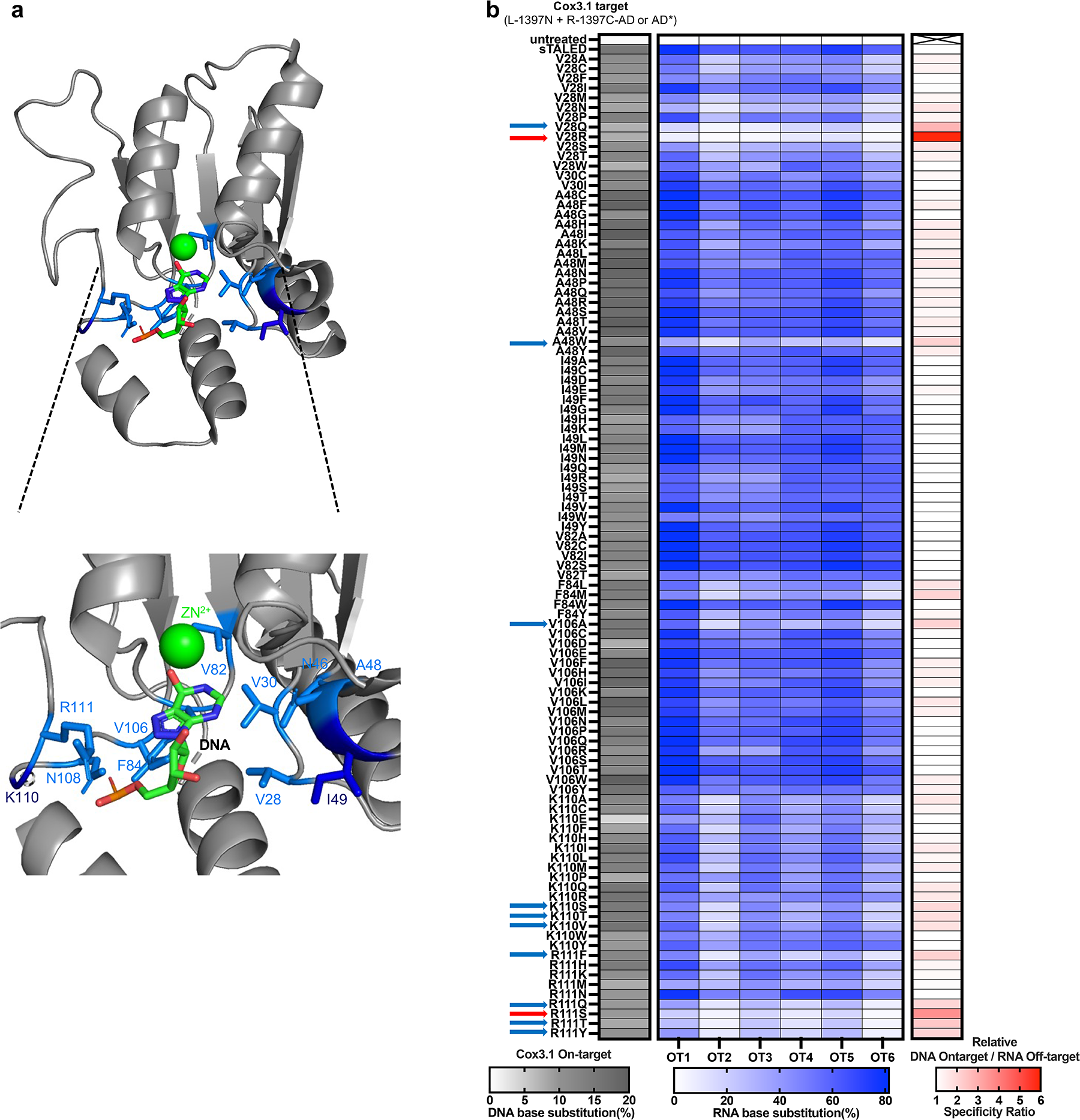
DNA on-target activity and RNA off-target activity of *COX3.1-*specific sTALED variants. (a) Structural representations of the TadA portion of ABE8e (Protein Data Bank (PDB) accession number 6VPC). The 11 amino-acid residues in close proximity with a DNA substrate are shown in sky blue or blue. Zinc ion is shown in green. 8-azanebularine (8AZ), a transition state analog for adenosine deamination reactions, is shown in the middle of the catalytic pore. PyMol (Methods) was used to create all graphical representations. (b) Heat map showing DNA on-target activity (left), RNA off-target activity (middle), and the relative ratio of DNA on-target editing frequencies to RNA off-target editing frequencies (right) of the 101 sTALED variants among the total of 209 sTALED variants (11X19) that retain mtDNA on-target activity. The relative ratio is normalized to that of the original sTALED, which has a value of 1. Data are shown as means from n = 3 biologically independent samples. Source data are provided as a Source Data file.

We next used targeted RNA amplicon sequencing to measure off-target editing frequencies of the 101 sTALEDs at six representative sites revealed by the transcriptome-wide sequencing analysis described above (Fig. 2b and Supplementary Fig. S6a). These representative sites were chosen because they were mutated with high frequencies that ranged from 60% to 80% by both of the *Cox3.1*-and *ND1*-specific TALEDs (Supplementary Fig. S6a). RNA off-target editing frequencies measured by targeted deep sequencing were in good agreement with those estimated by transcriptome-wide sequencing (Supplementary Fig. S6b and S6c). We chose a total of 12 TadA8e variants, which minimized RNA off-target editing efficiencies at the six representative sites, while retaining mtDNA on-target editing efficiencies (Fig. 2b).

### Transcriptome-wide off-target editing analysis

We next performed transcriptome-wide sequencing to investigate whether the 12 sTALED variants could avoid off-target editing at sites other than the six representative sites (Fig. 3). All of these variants reduced the number of off-target edits dramatically. For example, sTALED variants with V28R and R111S induced off-target edits at merely 852 and 829 sites, respectively, whereas the original sTALED and sTALED-V106W (sTALED with the V106W mutation in TadA8e) induced off-target edits at 96,559 and 81,156 sites, respectively (Fig. 3a and 3b). Thus, sTALED-V28R and -R111S avoided >99% of RNA off-target edits. Considering that 316 A-to-G edits were found in the untreated sample used as a negative control, which were likely caused by high-throughput sequencing errors, these results showed that these sTALED variants almost completely avoided RNA off-target editing. We also investigated whether these TadA8e mutations could reduce RNA off-target editing when incorporated into sTALEDs targeted to other mtDNA target sites (Fig. 3c-f and Supplementary Fig. S7). Targeted RNA amplicon sequencing showed that most of the *ND1*-and *ND6*-specific sTALED variants reduced RNA off-target editing frequencies at the six representative sites significantly (Supplementary Fig. S7c-f). We also measured mtDNA on-target editing frequencies via targeted deep sequencing (Supplementary Fig. S7a and S7b) and obtained the ratio of the DNA on-target editing frequency to the RNA off-target editing frequency (Supplementary Fig. S7g and S7h). Based on these results, we chose the following four TadA8e variants, V28Q, V28R, A48W, and R111S, which exhibited high mtDNA on-target activity and low RNA off-target activity, compared to the original sTALED and sTALED-V106W targeted to the *Cox3.1, ND1,* and *ND6* sites (Supplementary Fig. S7i). Whole transcriptome sequencing showed that *ND1*-and *ND6*-specific sTALED variants with each of these four variations reduced the number of RNA off-target edits drastically (Fig. 3), in line with the results shown above with the *Cox3.1*-specific sTALED variants. Again, both the *ND1*-and *ND6*-specific sTALED-V28R or -R111S variants were the most discriminatory, inducing the fewest RNA off-target edits (Fig. 3c-f). These new variants, when incorporated into sTALEDs with DddA_tox_ split at G1333, also reduced RNA off-target edits (Supplementary Fig. S8). We also tested whether V28R and R111S could be combined with each other and also with V106W to improve sTALEDs further: The resulting double or triple variants, however, were poorly active at the on-target sites (Supplementary Fig. S9).

**Figure 3.**
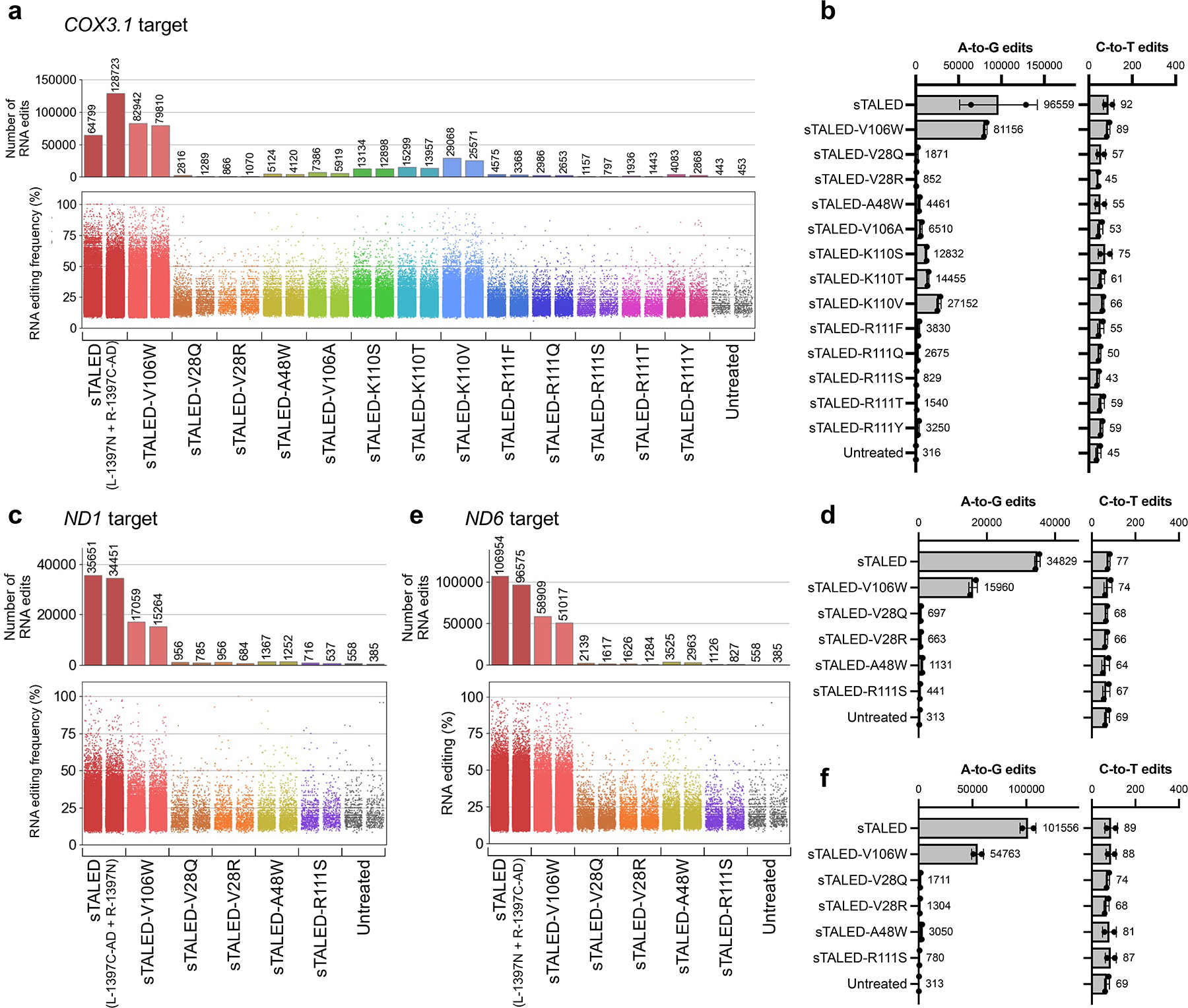
Transcriptome-wide sequencing to examine RNA off-target activity of sTALED variants. Bar graphs illustrating the total number of RNA edits found in HEK 293T cells that expressed sTALED or sTALED variants. Results from untreated cells are also shown. (a) Number of RNA edits (top panel) and editing frequencies of each RNA edit for the *Cox3.1*-specific sTALEDs shown as a dot (bottom panel). (b) A-to-G edits and C-to-T edits for the the *Cox3.1*-specific sTALEDs. (c) Number of RNA edits (top panel) and editing frequencies of each RNA edit for the *ND1*-specific sTALEDs shown as a dot (bottom panel). (d) A-to-G edits and C-to-T edits for the *ND1*-specific sTALEDs. (e) Number of RNA edits (top panel) and editing frequencies of each RNA edit for the *ND6*-specific sTALEDs shown as a dot (bottom panel). (f) A-to-G edits and C-to-T edits for the *ND6*-specific sTALEDs. Data are shown as means from n = 2 biologically independent samples. Source data are provided as a Source Data file.

We next analyzed whole transcriptome sequencing data to investigate whether sTALEDs could induce RNA off-target edits in the mitochondrial transcriptome and whether the two variants could reduce or avoid them. A-to-I edits were not induced at high frequencies in mitochondrial RNAs by the three sTALEDs targeted to the *Cox3.1*, *ND1*, and *ND6* sites. We were, however, able to find hundreds of low-frequency (<10%) off-target A-to-I edits in the transcriptome (Supplementary Fig. S10a). Thus, at the mtRNA-wide level, the three original sTALEDs induced random A-to-I editing with average frequencies that ranged from 0.051% to 0.11% (0.076 ± 0.018%), 2.3∼5.0-fold higher than that observed with the untreated control (0.022%). sTALED-V28R and -R111S targeted to the same three sites reduced RNA off-target editing significantly, inducing average frequencies of 0.027±0.008% and 0.033±0.003%, respectively (Supplementary Fig. S10b).

### Engineered TALEDs reduce bystander and off-target DNA editing

We reasoned that sTALED variants with site-specific mutations at the TadA8e substrate-binding site could also reduce bystander editing at the target site and off-target editing in the mitochondrial genome, because these mutations could potentially fine-tune adenine deaminase activity for DNA substrates in addition to reducing activity for RNA substrates. First, we examined base editing frequencies at each nucleotide position. Both sTALED-V28R and -R111S variants specific to each of a total of eight mtDNA sites, including the *Cox3.1, ND1,* and *ND6* sites, induced A-to-G edits in a narrower window than did the wild-type sTALEDs or sTALED-V106W variants targeted to the same sites (Fig. 4 and Supplementary Fig. S11). For example, the wild-type sTALED and sTALED-V106W targeted to the *Cox3.1* site induced A-to-G edits at multiple positions, with frequencies of >1.1%, not only in the spacer region between the two TALE-binding sites but also in the TALE-binding sites. In contrast, the *Cox3.1-* specific sTALED-V28R induced an A-to-G edit at a single position in the spacer region (Fig. 4a). The other sTALEDs and sTALED variants showed a similar trend. In summary, the V28R and R111S variants were as efficient as the original sTALEDs at the middle position (A10 in Supplementary Fig. S11g), but, unlike the original sTALEDs, rarely induced A-to-G edits at positions several nucleotides upstream (A1 to A6) or downstream (A15 to A20). In particular, the V28R variants, targeted to 4 sites (*ND1, ND6, ND1-3*, and *ND5-2*) among the total of eight mtDNA sites, induced A-to-G edits more efficiently at the peak editing position (boxed in Fig. 4a-c and Supplementary Fig. S11) in each target site, compared to the original sTALEDs. These results suggest that the narrowed editing window observed with the two new TALED variants is not necessarily a byproduct of their poor on-target activity.

**Figure 4.**
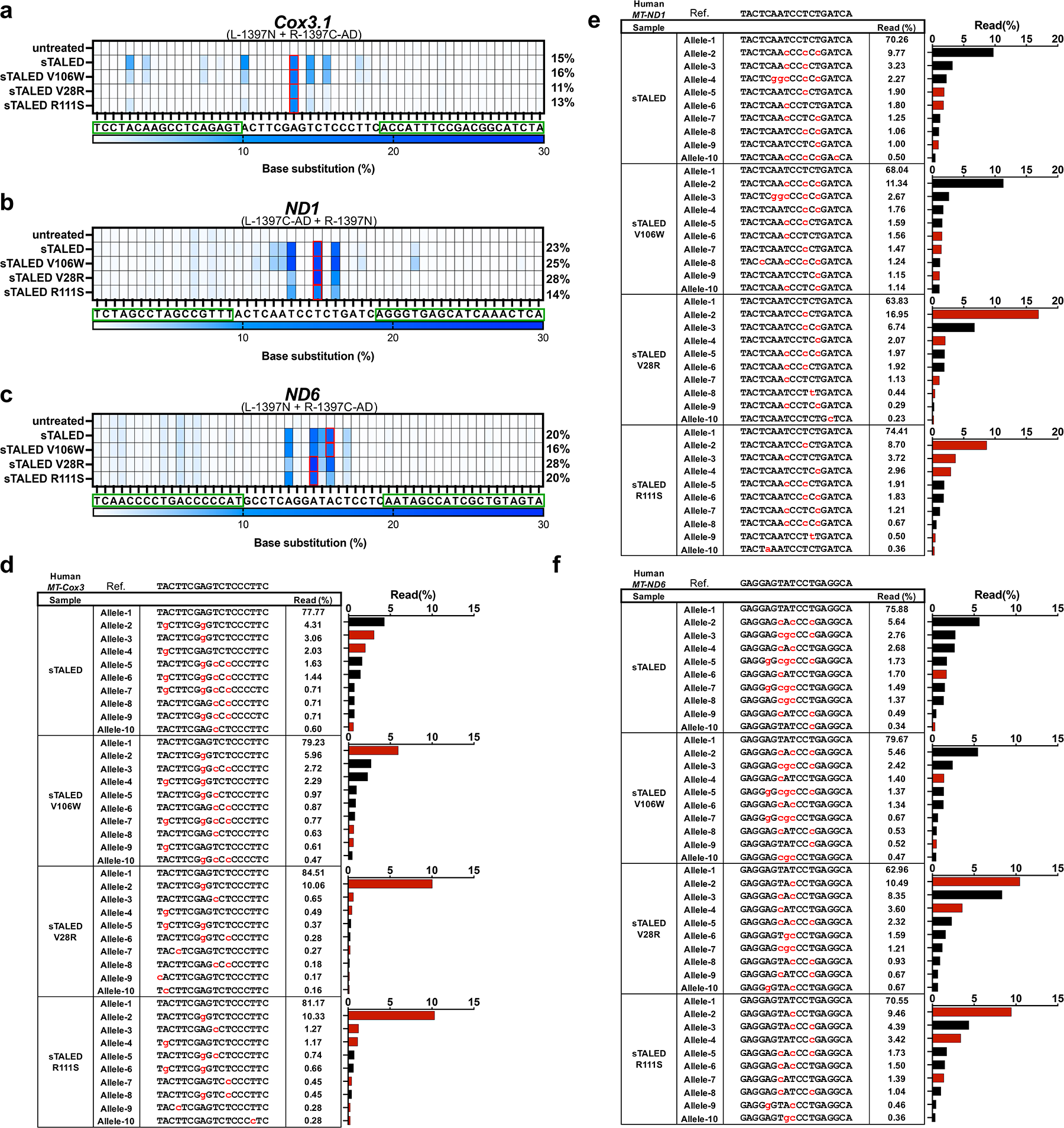
Editing patterns induced by *Cox3.1*-*, ND1*-, and *ND6*-specific sTALED variants. Heat maps depicting A-to-G conversions caused by sTALED or sTALED variants targeted to the *Cox3.1* site (a), *ND1* site (b), and *ND6* site (c) in HEK 293T cells. TALE-binding sites are indicated by the green boxes. Data are shown as means from n = 2 biologically independent samples. (d) Analysis of *Cox3.1* alleles summarized in (a). (e) Analysis of *ND1* alleles summarized in (b). (f) Analysis of *ND6* alleles summarized in (c). (d-f) The spacer sequence is shown, with A-to-G nucleotide changes in red (left). Bar graphs displaying the frequency of each allele are shown on the right. Bars representing alleles with only single-nucleotide conversions are shown in red. Only the top 10 sequence reads among hundreds of edited sequences are shown. Source data are provided as a Source Data file.

Next, we compared the frequencies of edited alleles induced by these sTALEDs targeted to the *Cox3.1, ND1,* and *ND6* sites (Fig. 4d-f). Most of the mutant alleles induced by the original sTALEDs or sTALED-V106W variants contained multiple-base edits rather than single-base edits. In sharp contrast, sTALED-V28R and -R111S induced mutant alleles with single-base substitutions much more frequently than multiple-base substitutions. Thus, the most abundant alleles with a single A-to-I edit in the middle of the spacer region were induced with low frequencies of 3.06% (*Cox3.1*), 1.90% (*ND1*), and 1.70% (*ND6*) by the original sTALEDs, whereas the same alleles were induced with high frequencies of 10.1% (*Cox3.1*), 17.0% (*ND1*), and 10.5% (*ND6*) by the sTALED-V28R variants (Fig. 4d-f). This result has an important implication for the use of TALEDs in disease modeling. TALEDs inducing single-base substitutions with no or few bystander edits are desired, because the vast majority of pathogenic mtDNA mutations, responsible for mitochondrial genetic disorders, are single-nucleotide variations rather than multiple-nucleotide variations.

We next carried out whole mitochondrial genome sequencing to assess and compare off-target effects of the wild-type sTALEDs and sTALED variants at day 4 post-transfection (Fig. 5). In this analysis, we excluded several naturally-occurring variations, with a heteroplasmy fraction of ≥ 10%, found even in the untreated sample and off-target edits observed in TALED-treated or untreated samples with low frequencies of ≤ 0.1%, which were likely to arise from high-throughput sequencing errors. The wild-type sTALEDs targeted to the three mtDNA sites induced off-target mutations with average frequencies of mitochondrial genome-wide off-target editing that ranged from 0.0024% to 0.0066%, 4∼10-fold higher than that observed in the untreated control (0.0006%). In contrast, sTALED-V28R and R111S induced off-target edits with average frequencies of < 0.0006%, similar to the baseline frequency observed in the negative control (Fig. 5b). Furthermore, the number of off-target edits induced in the mitochondrial genome was also reduced by the sTALED variants. Thus, the *ND1*-specific, wild-type sTALED caused A-to-I off-target edits at 108 sites in human mtDNA with frequencies of > 0.1%, whereas sTALED-V28R and -R111S induced off-target edits at 14 and 17 sites, respectively, similar to the baseline number (that is, 24) seen in the untreated sample (Fig. 5a). We also performed whole mitochondrial genome sequencing at day 2 post-transfection (Supplementary Fig. S12), when RNA off-target edits were induced at the highest level. The original sTALEDs targeted to the three sites induced off-target edits with average frequencies that ranged from 0.0071% to 0.0090%, 11∼14-fold higher than that observed in the untreated control (0.0006%). However, the V28R and R111S variants did not induce off-target edits, compared to the negative control (Supplementary Fig. S12b).

**Figure 5.**
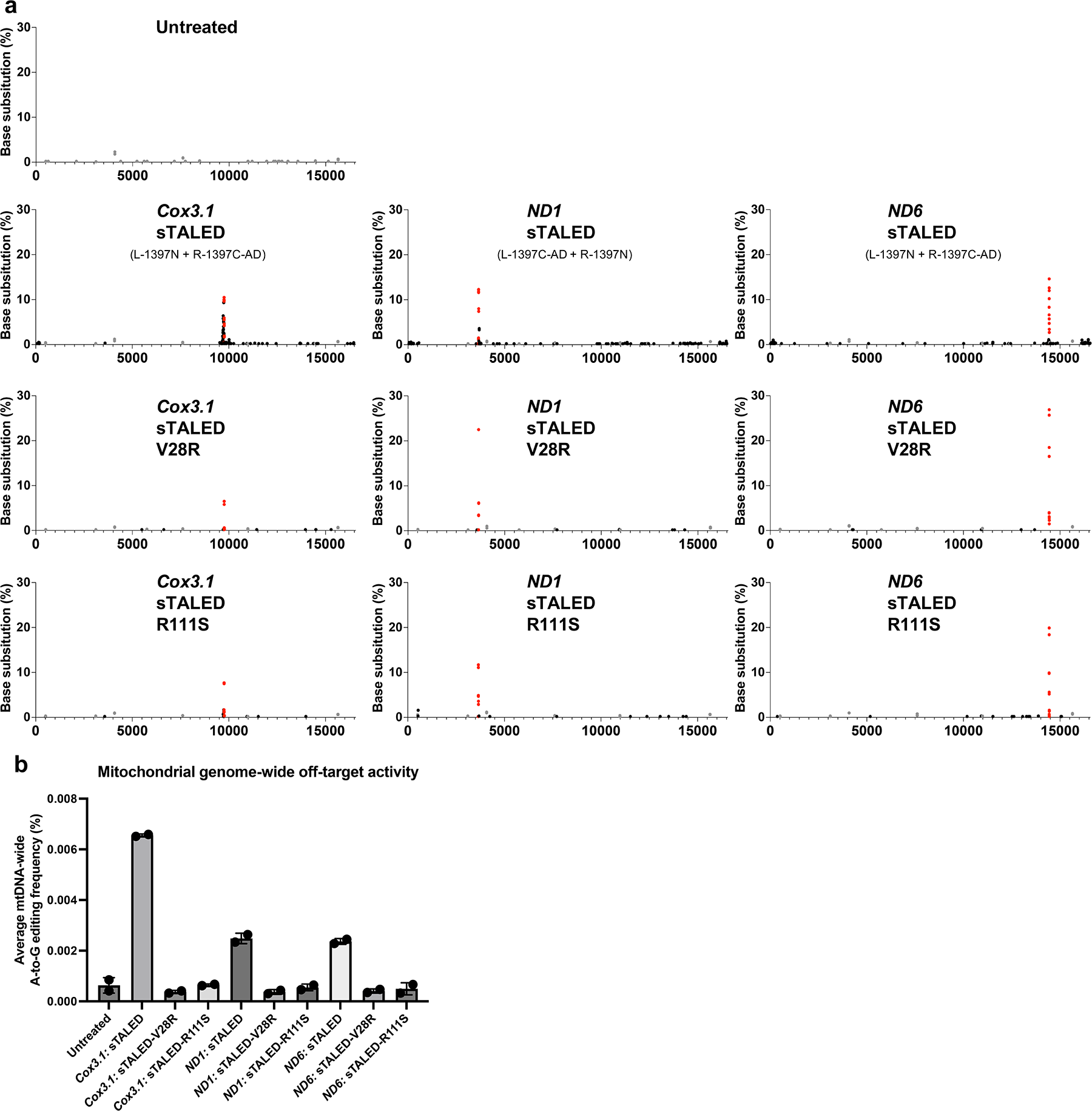
Whole mitochondrial genome sequencing to examine off-target effects of sTALED and sTALED variants. (a) Plots indicate the positions of on-target and off-target edits across the mitochondrial genome at day 4 post-transfection. Dots represent A-to-G substitutions with frequencies of > 1%. Red, black, and gray dots indicate on-target (and bystander) edits, off-target edits, and naturally-occurring single-nucleotide variations (SNVs), respectively. Nucleotide positions in the human mitochondrial genome are represented on the X axis. All data points from nL=L2 biologically independent experiments are shown. (b) The average frequencies of genome-wide off-target edits induced by wild-type sTALED and sTALED variants targeted to the indicated sites. Error bars are s.e.m. for nL=L2 biologically independent samples. Source data are provided as a Source Data file.

### Time course measurements of DNA on-target and RNA off-target editing frequencies

We next investigated whether on-target mutations induced by various forms of sTALEDs were stably maintained and how long RNA off-target variations persisted over time (Supplementary Fig. S13). We measured, using targeted deep sequencing, DNA on-target editing frequencies at the three mtDNA sites (Supplementary Fig. S13a-c) and RNA off-target editing frequencies at the six representative sites (Supplementary Fig. S13d-g), co-edited by sTALEDs targeted to the three sites, at various time points. As expected, RNA edits were heavily induced by sTALEDs and sTALED-V106W variants at day 1 and 2 post-transfection but almost completely disappeared by day 8 post-transfection. This result is not surprising because sTALED expression plasmids were transiently, rather than stably, transfected into cells. The new sTALED variants reduced RNA off-target editing frequencies by several fold, compared to sTALEDs or the sTALED-V106W variants, even at day 1 and 2 post-transfection (Supplementary Fig. S13g). We also found that mtDNA on-target edits induced by the sTALEDs and sTALED-V106W variants were not stably maintained. Thus, the frequencies of mtDNA on-target edits induced by the sTALEDs and sTALED-V106W variants dropped significantly over time. For example, mtDNA on-target edits were induced by the *ND6*-specific sTALED and sTALED-V106W with high frequencies of up to 46% and 39%, respectively, at day 1 and 2 post-transfection but with low frequencies of < 9% at day 8 or later (Supplementary Fig. S13c). In contrast, the frequencies of on-target edits induced by sTALED-V28R and -R111S increased or were more stably maintained over time. As a result, the editing frequencies observed for our new sTALED variants were lower at day 1 and 2 post-transfection but were higher at day 8 or later than those induced by previous versions of sTALEDs (Supplementary Fig. S13a-c).

Based on these results, we reasoned that sTALEDs and sTALED-V106W variants were cytotoxic, because they induced too many RNA off-target edits with high frequencies, and that mtDNA-edited cells could not divide or survive over time. sTALED-V28R and -R111S variants could be tolerated, possibly because they avoided RNA off-target editing or did not induce too many bystander edits at the target site and off-target mutations in the mitochondrial genome. We performed cell proliferation assays (Supplementary Fig. S14a) and found that, indeed, the sTALEDs and sTALED-V106W variants were cytotoxic, reducing cell viability significantly, compared to the negative control (pEGFP transfection) (Supplementary Fig. S14b). sTALED-V28R and -R111S were tolerated much better, such that cell viability was not reduced at day 4 post-transfection (Supplementary Fig. S14c).

### TadA8e-V28R and -R111S incorporated into CRISPR RNA-guided ABEs

We investigated whether the V28R and R111S mutations in TadA8e could also reduce bystander editing and RNA off-target editing induced by CRISPR RNA-guided ABEs, widely used for nuclear DNA editing (Supplementary Fig. S15). First, we measured on-target and bystander editing frequencies at a total of eight target sites. Both ABE8e-V28R and -R111S were as efficient as ABE8e and ABE8e-V106W (ABE8eW) at these target sites with an average editing frequency of 33% (V28R) and 41% (R111S) (Supplementary Fig. S15b and S15c). Interestingly, the V28R and R111S variants exhibited a narrowed editing window with maximum editing at position A5 (35% and 45%, respectively) in the proto-spacer region and minimal bystander editing at A3 (1.0% and 2.9%, respectively) and A10 (0.73% and 0.65%, respectively) (Supplementary Fig. S15d and S15e). In contrast, ABE8e and ABE8eW showed a broader editing window with maximum editing at positions A5 (46% and 48%, respectively) and A7 (39% and 42%, respectively) and substantial bystander editing at A3 (24% and 19%, respectively) and A10 (19% and 13%, respectively).

We next used targeted RNA amplicon sequencing to assess RNA off-target editing activities. ABE8e-V28R and -R111S reduced average RNA off-target editing frequencies measured at six representative RNA off-target sites by 9.9-fold and 4.7-fold, respectively, compared to ABE8e, and 6.3-fold and 3.1-fold, compared to ABE8eW (Supplementary Fig. S15f and S15g). Furthermore, whole transcriptome sequencing showed that ABE8e-V28R and -R111S reduced the number of RNA off-target edits substantially, compared to ABE8e and ABE8eW (Supplementary Fig. S16a and S16b). These results show that the two new TadA8e variations, V28R and R111S, can also improve ABE8e by reducing RNA off-target edits and bystander edits.

### Mitochondrial DNA adenine base editing in mice

In parallel with engineering TALEDs to reduce RNA off-target effects, we attempted to create animal models with pathogenic mtDNA mutations by using custom-designed TALEDs targeted to two mitochondrial genes, *mtATP6* and *mtRnr1*. Both sTALED targeted to *mtATP6* and mTALED targeted to *mtRnr1* (Fig. 6a) achieved efficient A-to-G edits at the two target sites with high frequencies of up to 22% in NIH3T3 cells and up to 19% in B16F10 cells (Supplementary Fig. S17). We next injected in vitro transcribed mRNAs encoding mTALED or sTALED into mouse one-cell embryos. Unexpectedly, more than 40% of the TALED-injected embryos failed to form blastocysts and growth-arrested at the 2-cell or 4-cell stage. A-to-G edits were observed in most of the resulting arrested embryos with an average frequency of 4% (*mtRnr1*) or 26% (*mtATP6*) (Fig. 6b). In sharp contrast, adenine base edits were not detectably induced in 29 of 30 blastocysts injected with the *mtATP6*-specific sTALED or in 18 of 19 blastocysts injected with the *mtRnr1*-specific mTALED. A low-dosage (10 ng/uL) injection of sTALED into embryos, however, allowed blastocyst development in 11 of 14 embryos. But adenine editing frequencies were low (< 7%) in the resulting blastocysts (Fig. 6c). These results suggest that mouse embryos failed to divide due to cytotoxicity of the TALEDs, which was caused by the presence of TadA8e in mTALED or in one subunit (R-1397C-AD) of a sTALED pair (L-1397N + R-1397C-AD). Indeed, none of the eight embryos injected with the left-side subunit, L-1397N, were growth-arrested and all divided normally to form blastocysts, whereas 7 of 17 embryos injected with the right-side subunit containing TadA8e, R-1397-AD, were arrested at the 2-cell or 4-cell stage (Fig. 6c).

**Figure 6.**
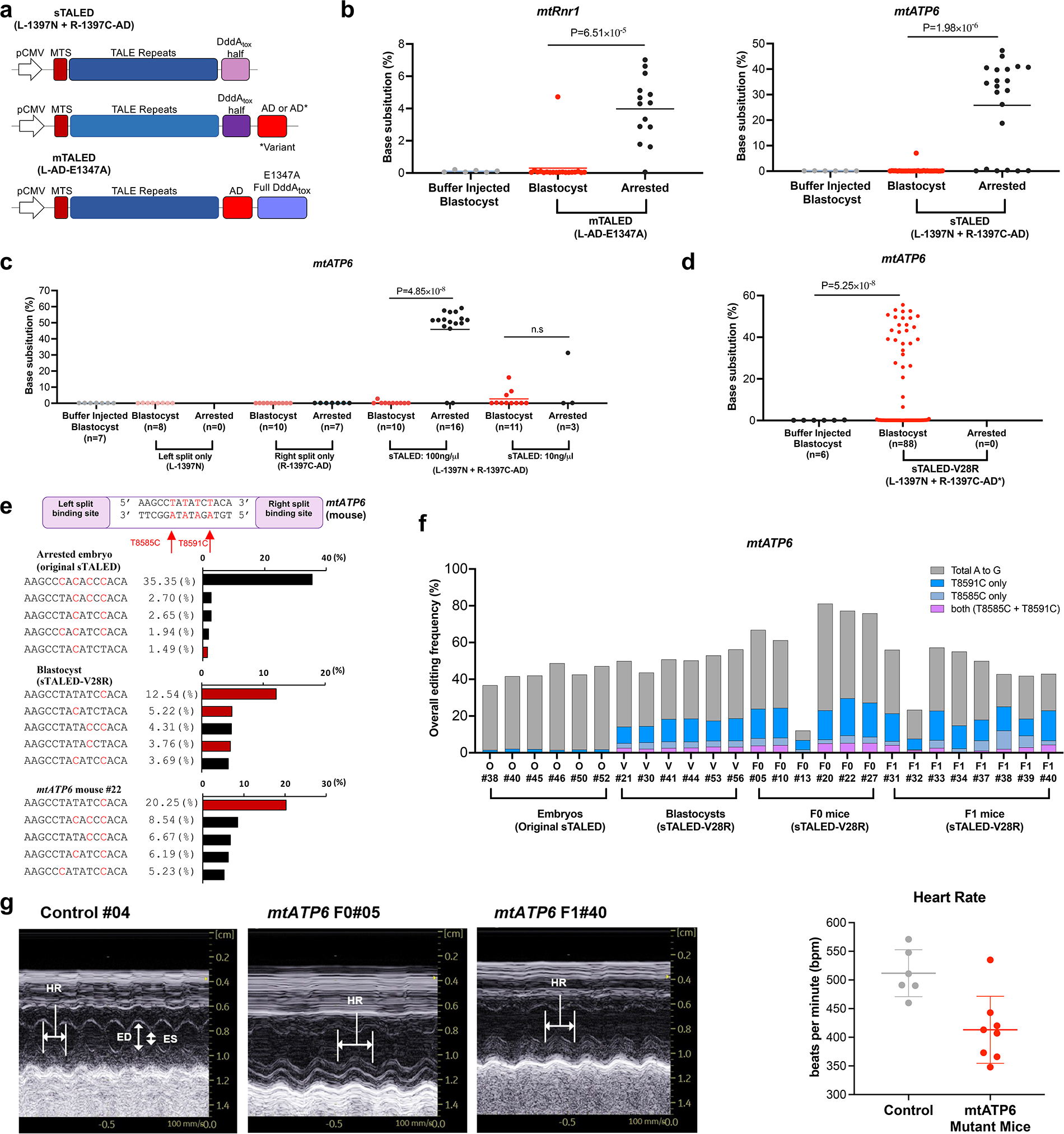
Mouse mitochondrial DNA mutations induced by the sTALED-V28R variant. (a) Architecture of the sTALED, sTALED variant, and mTALED constructs used in mouse experiments. AD, TadA8e adenine deaminase; AD*, TadA8e adenine deaminase variant; MTS, mitochondrial targeting sequence. Light purple indicates the N-terminal halves of DddA_tox_ and dark purple indicates the C-terminal halves of DddA_tox_. (b) Editing frequencies in TALED-injected embryos. mTALED was targeted to the mouse *mRNR1* site and sTALED was targeted to the *mtATP6* site. (c) Adenine base editing frequencies in blastocysts and arrested embryos. Each subunit of the *mtATP6*-specific sTALED was injected into mouse embryos. The sTALED pair was also injected at a low (10 ng/uL) or high (100 ng/uL) dosage. (d) Adenine base editing induced by the sTALED-V28R variant and associated embryo development or growth arrest. (e) *mtATP6* mutant alleles induced in the sTALED-treated embryos and mouse. (Upper panel) The target spacer region is indicated, with A-to-G nucleotide changes shown in red. Red arrows indicate two nucleotide positions with pathogenic mutations. (Lower panel) Frequencies (%) of *mtATP6* mutant alleles. The red or black bars represent alleles with single-nucleotide or multiple-nucleotide conversions, respectively. (f) Distribution of *mtATP6* mutant alleles induced in embryos and mice. Mutant alleles are divided into four groups: T8585C only, T8591C only, Both (= T8585C + T8591C), and alleles with bystander edits. (g) Echocardiography (left) and heart rates (bpm) (right) of *mtATP6*-edited and control mice. In the echocardiograms, HR measurements from six sites based on ED-ED* and ES-ES* were averaged for each mouse to measure heart rate in bpm. ES=end systole, ED=end diastole, HR=heart rate. Source data are provided as a Source Data file.

Having confirmed that TadA8e in TALEDs caused growth arrest in mouse embryos, we reasoned that our TALED variants with minimum RNA off-target editing activity could be used to create mouse models with pathogenic mtDNA mutations. Unlike the sTALEDs described above, the *mtATP6*-specific sTALED-V28R did not cause growth arrest: all 88 injected embryos gave rise to blastocysts (Fig. 6d). Furthermore, 28 of 88 blastocysts contained A-to-G base edits in the mitochondrial gene with editing frequencies ranging from 6% to 54% (39%, on average). We also noted that sTALED-V28R induced mutant alleles with single-base edits preferentially over those with multiple-base edits in blastocysts, whereas the original sTALED induced those with multiple-base edits almost exclusively in arrested embryos (Supplementary Fig. S18), suggesting that the V28R variant could reduce or avoid bystander edits at the target site.

We next implanted sTALED-V28R-injected embryos into surrogate mothers and obtained F0 pups with mtDNA mutations (Fig. 6e-g). These mice carried two mutant alleles without bystander edits (m.T8585C and m.T8591C, corresponding to m.T9185C and m.T9191C, respectively, in the human mitochondrial genome)^40^, associated with Leigh disease, at a combined frequency of 22.5±3.3%. In particular, one F0 mouse (#22) carried the m.T8591C allele without bystander edits at a frequency of 20.3% (Fig. 6e and Supplementary Fig. S19a). This single base-edited allele was induced at a low frequency of 1.30±0.08% in mouse embryos injected with the original sTALED. These two pathogenic alleles were transmitted maternally to the next (F1) generation (Fig. 6e-g and Supplementary Fig. S19b). Notably, these F0 and F1 mutant mice showed reduced heart rates (410±21 beats per minute (bpm)), compared to wild-type siblings (510±17 bpm), a phenotype observed previously in a nuclear gene-disrupted mouse model (*Ndufs4* -/-) of Leigh syndrome ^41^ (Fig. 6g and Supplementary Fig. S20). No off-target mutations were detectably induced at a total of 18 candidate sites with high sequence homology in the nuclear genome (Supplementary Fig. S21). High-throughput mitochondrial genome-wide sequencing also showed that no off-target mutations were detectably induced in our mutant mice (Supplementary Fig. S22).

## DISCUSSION

Adenine and cytosine base editors enabling targeted base substitutions in mitochondrial DNA are broadly useful in research and medicine^10–13, 42–47^, but are limited by off-target editing in the mitochondrial genome and bystander editing at the target site. In addition to these limitations, here, we showed that adenine editing TALEDs but not cytosine editing DdCBEs could induce tens of thousands of unwanted RNA off-target edits in human cells. Although most of these RNA off-target edits disappeared several days after transfection, TALEDs reduced cell viability and caused developmental arrest of mouse embryos. To overcome these limitations of TALEDs, we constructed and tested a total of 209 TALED variants in which each of the 11 amino-acid residues near the substrate-binding pocket in the TadA protein was replaced with the other 19 amino-acid residues. As a result, we were able to develop TALED variants with fine-tuned deaminase activity, avoiding > 99% of RNA off-target edits and minimizing off-target edits and bystander edits. Interestingly, our engineered TALEDs induced on-target A-to-G edits less efficiently at day 1 or 2 post-transfection but more efficiently at day 4 or later than did the original sTALED and sTALED-V106W variant (Supplementary Fig. S13). In fact, most of the on-target edits induced by sTALEDs or sTALEDs-V106W disappeared by day 8 post-transfection, whereas those induced by our sTALED variants were more stably maintained over time. This result suggests that sTALEDs and sTALEDs-V106W were cytotoxic, due to RNA off-target activity, and that our engineered sTALED variants were tolerated, because they reduced RNA off-target editing. In line with this result in cultured human cells, TALED-injected mouse embryos failed to divide and form blastocysts. Fortunately, we were able to use sTALED-V28R to obtain mtDNA-edited mice with pathogenic mutations associated with Leigh syndrome.

We also found that the V28R and R111S variations in TadA8e could be introduced into CRISPR RNA-guided ABEs to avoid RNA off-target edits and to reduce bystander edits. Further studies are needed to determine whether these variations are better than or synergistic with several other variations in TadA8e that had been shown to fine-tune RNA-guided ABE activity. We note that some of the previously-known variations, including V106W^35^, V106G^34^, and N108Q^29, 38^, were included in our initial library of 209 TALED variants. sTALED variants with V106G or N108Q, however, were poorly efficient and excluded from later experiments. Various V28 mutations other than V28R have also been incorporated into ABE8e ^38, 48, 49^. It is possible that these variations are compatible with CRISPR RNA-guided ABEs targeting nuclear DNA but not with TALEDs targeting mtDNA.

ABEs have been successfully used for base editing in animal models without apparent cytotoxicity^38, 50–54^. We speculated that MTS-induced mitochondrial translocation of TALEDs is less efficient or much slower than is nuclear localization signal (NLS)-induced nuclear translocation of ABEs. Indeed, we found that ABE8e containing MTS (derived from COX8a or SOD2) rather than NLS induced RNA off-target edits much more frequently at the six representative sites than did the original ABE8e containing NLS (Supplementary Fig. S23a). Furthermore, cell proliferation assays showed that these variants were more cytotoxic than the original ABE8e (Supplementary Fig. S23b). In a reciprocal experiment, sTALED variants, in which MTS was replaced by NLS, exhibited significantly reduced RNA off-target editing (Supplementary Fig. S23c). These results suggest that TadA8e fused to MTS but not to NLS is problematic.

TALEDs induced heteroplasmic editing rather than homoplasmic (that is, 100%) editing in mtDNA both in cultured mammalian cells and one-cell stage mouse embryos. Heteroplasmic editing may be sufficient for phenotypic effects in some cases. If not, homoplasmic editing can be achieved by stable transfection or long-term expression of TALEDs. In fact, we were able to obtain homoplasmic editing in plastid DNA in whole plants using stable transfection of TALEDs via agrobacterium^42^ and in mtDNA in mammalian cells^21^ using adeno-associated virus expressing monomeric DdCBEs. It remains to be tested whether our TALED variants with reduced RNA off-target activity are more suitable for long-term expression.

## ACKNOWLEDGEMENTS

This work was supported by the Institute for Basic Science grant (IBS-R021-D1), the Korea Research Institute of Bioscience and Biotechnology (KRIBB) Research Initiative Program (KGM1012311), the National Research Foundation of Korea (NRF) grants funded by the Korea government (MSIT) (RS-2023-00261905, RS-2023-00210965, RS-2023-00260462, and RS-2023-00260462), and the Korea Institute of Science and Technology (KIST) Institutional Program (2E32161).

## AUTHOR CONTRIBUTIONS

J.-S.K. supervised the research. S.-I.C. and J.-S.K. conceived the research. S.-I.C., K.L., S.L., H.L., and J.-S.K. design the study. S.-I.C., K.L., H.S., S.L., and H.L. performed and analyzed the main experiments. J.L., A.K., J.M.L., Y.G.M., E.C., S.H., S.-M.C., J.K., S.K., E.-K.K., K.-H.N., Y.O., and M.C. performed the experiments. S.-I.C., K.L., S.L., H.L., and J.-S.K. wrote the manuscript.

## AUTHOR INFORMATION

J.-S.K. is a co-founder of and holds stock in ToolGen, Inc., Edgene, Inc., and GreenGene, Inc. S.-I.C., K.L., J.L., S.L., H.L., and J.-S.K. have filed patent applications related to this work. The other authors declare no competing interests. Correspondence and requests for materials should be addressed to J.-S.K.

## METHODS

### PyMOL analysis

The ABE8e (PDB accession number 6VPC^39^) structure was downloaded from PDB and visualized with PyMOL v.2.5.4. Some elements including Cas9, single guide RNA, and double-stranded DNA were excluded from the PDB file; 8AZ, a transition state analog for adenosine deamination reactions, and the TadA monomer were retained. We selected 11 residues (V28, V30, N46, A48, I49, V82, F84, V106, N108, K110, and R111) that were near 8AZ, including residues previously known to contact DNA.

### Plasmid construction

To construct plasmids encoding DdCBEs, sTALEDs, mTALEDs, and dTALED that target specific sites, we began with plasmids containing a stuffer, which is a sequence between two restriction enzyme sites that is helpful when separating fragments during gel electrophoresis. The architectures of the plasmids are as follows: sTALED plasmids (p3s-stuffer-DddA_tox_ half (1397C)-AD and p3s-stuffer-DddA_tox_ half (1397N) or p3s-stuffer-DddA_tox_ half (1333C)-AD and p3s-stuffer-DddA_tox_ half (1333N)); mTALED plasmids (p3s-stuffer-E1347A DddA_tox_ full-AD); dTALED plasmids (p3s-stuffer-AD and p3s-stuffer-19 E1347A DddA_tox_ full). These plasmids were obtained from Addgene (DdCBE, #187168, #187171, #187173, #178174; sTALED, #187167, #187169, #187170, #187172; mTALED, #187163, #187166; dTALED, #187164, #187165). Plasmids encoding DdCBEs, sTALEDs, and mTALEDs targeting specific sites were constructed by inserting custom-designed TALE array sequences, similar to what was previously done in the construction of a TALEN library^55^. To remove the stuffer sequences, each plasmid digested with BsaI or BsmbI (NEB), and inserts, including a custom-designed TALE array sequence synthesized by IDT, were inserted into the digested vector using a HiFi DNA assembly kit (NEB). Alternatively, as in the previous work, the desired TALED construct was generated by digesting the master vector with BsaI or BsmbI to cleave the site in the stuffer and then assembling six TALE arrays in that position using the Golden Gate method.

The plasmids encoding ABE8e and ABE8e-V106W were obtained from Addgene (#138489 and #138495, respectively). The plasmid encoding ABE8e was modified to incorporate mutations in the adenine deaminase domain (V28R or R111S) using a Q5 Site-Directed Mutagenesis Kit (NEB).

### Cell culture and transfection

HEK 293T (CRL-11268, ATCC) cells were grown in DMEM (Welgene) with 10% fetal bovine serum (Welgene) and 1% antibiotic-antimycotic solution (Welgene). NIH3T3 (CRL-1658, ATCC) and B16F10 (CRL-6475, ATCC) cells were grown in DMEM supplemented with 10% (v/v) bovine calf serum (Gibco) for NIH3T3 cells or 10% fetal bovine serum (Gibco) for B16F10 cells in the absence of any antibiotics. Cell lines were maintained at 37°C with 5% CO_2_ and were passaged before reaching 90% confluency, with timing depending on the doubling period of the specific cell line.

For HEK 293T cell transfections, cells were seeded onto 48-well plates (Corning) at a density of 7.5 × 10^4^ cells per well prior to transfection. The cells were transfected with plasmids (total 1 ug) using 1.5 uL of Lipofectamine 2000 (Invitrogen) after 24 h. For delivery of a sTALED pair, a DdCBE pair, or a CRISPR-Base Editor with sgRNA, the total amount of plasmid was 1 ug (500 ng each). When a single construct was used, the amount of transfected plasmid was 500 ng. After 96 h, the transfected cells were harvested. For NIH3T3 and B16F10 cell transfections, cells were seeded in 24-well cell culture plates (SPL, Seoul, Korea) at a density of 1 × 10^5^ cells per well, 18–24 h before transfection. Lipofection using Lipofectamine 3000 (Invitrogen) was performed with 500 ng of each sTALED-encoding plasmid to make up 1000 ng of total plasmid DNA. For mTALED, 500 ng of plasmid was used. Cells were harvested 3 days after transfection.

### mRNA preparation

We used TALED-encoding plasmids from a previous study^11^. TALE binding sequences were replaced using the Golden Gate cloning method^55^ for each DNA site. Template DNAs were prepared by polymerase chain reaction (PCR) using Q5 High-Fidelity DNA Polymerase (NEB) with the following primers: F: 5′-CATCAATGGGCGTGGATAG-3′ and R: 5′-GACACCTACTCAGACAATGC-3′. mRNAs were synthesized using an in vitro RNA transcription kit (mMESSAGE mMACHINE T7 Ultra kit, Ambion), and purified with a MEGAclear kit (Ambion). All procedures were conducted according to the manufacturer’s instructions.

### Animals

Hormone-injected wild-type C57BL/6N females were mated to wild-type C57BL/6N males to generate embryos for collection, and female mice from the ICR strain were used as surrogate mothers. The mice were maintained in a specific pathogen-free facility under a 12-h dark-light cycle with constant temperature (20–26 °C) and humidity (40–60%).

### Microinjection of mouse zygotes

In preparation for zygote microinjection, 4-week-old C57BL/6N female mice (Taconic) were superovulated by injecting them with pregnant mare serum gonadotropin (5 IU, Prospec) and human chorionic gonadotropin hormone (5 IU, Sigma-Aldrich) intraperitoneally at 48-h intervals. For microinjection of sTALED-encoding mRNAs, split left and right TALED-encoding mRNAs (500 ng/μL each) were mixed and diluted in diethyl pyrocarbonate (DEPC)-treated injection buffer (0.25 mM ethylenediaminetetraacetic acid [EDTA], 10 mM Tris; pH 7.4). mTALED-encoding mRNA (500 ng/μL) was directly diluted in DEPC-treated injection buffer. Mixtures were then microinjected into the cytoplasm of zygote stage embryos, using a Nikon ECLIPSE Ti micromanipulator and a FemtoJet 4i microinjector (Eppendorf). The embryos were then cultured in microdrops of KSOM+AA (Millipore) at 37 °C for 2 days in a humidified atmosphere containing 5% CO_2_. Two-cell stage embryos were implanted into the oviducts of 0.5-dpc pseudo-pregnant surrogate mothers.

### Genotyping embryos and mouse cell lines

DNA was extracted from various stages of embryos by incubating them in lysis buffer (25 mM sodium hydroxide, 0.2 mM EDTA; pH 10) at 95 °C for 20 min, after which the pH was adjusted to 7.4 using 4-(2-hydroxyethyl)-1-piperazineethanesulfonic acid (HEPES, free acids without pH adjustment) at a final concentration of 50 mM. For mouse cell lines (NIH3T3, B16F10), genomic DNA was extracted for PCR genotyping using DNeasy Blood & Tissue Kits (Qiagen), and subjected to targeted deep sequencing.

### Transcriptome sequencing

Total RNA was isolated using a NucleoSpin RNA Kit (MN #Macherey-Nagel) according to the manufacturer’s instructions at 48 h or 96 h post-transfection. RNA libraries were prepared using a TruSeq Stranded Total RNA Library Prep Gold kit (Illumina). RNA library quality was assessed using a 2200 TapeStation with a D1000 ScreenTape system (Agilent). Total RNA sequencing was performed using a NovaSeq 6000 Sequencer (Illumina) at Macrogen with paired-end sequencing systems (2 x 100bp).

### RNA variant calling

To analyze next-generation sequencing (NGS) data from RNA sequencing, we referred to a RNA variant-calling pipeline previously used for RNA off-target analysis^32, 34^ of CRISPR DNA base editors. Briefly, we aligned the fastq sequencing reads to the hg38 human reference genome (GRCh38, release v105) using STAR aligner (v.2.7.10a). The resulting BAM files were processed with GATK (v.4.2.4.1) MarkDuplicates, BaseRecalibrator, and ApplyBQSR. RNA base-editing variants were called using GATK HaplotypeCaller. RNA variant loci were compared to those of control samples and filtered based on the following criteria: (1) loci with a read depth of at least 10 were retained; (2) loci with a variant count of at least 2 were retained; (3) loci also present in the control sample were removed; and (4) undeterminable loci due to insufficient sequencing depth in the control sample were excluded. For the replicate-1 experimental sets, untreated replicate-2 was used as the control for filtering, and for the replicate-2 experimental sets, untreated replicate-1 was used as the control. A-to-G edits were counted as the number of RNA variant loci with A-to-G edits on the positive strand or T-to-C edits on the negative strand. C-to-T edits were counted as the number of RNA variant loci with C-to-T edits on the positive strand or G-to-A edits on the negative strand.

To further analyze the mitochondrial transcriptome from RNA sequencing, we extracted the base sequences and read counts for each nucleotide position from the processed BAM file using Bam-readcount ^56^. Positions with a read depth <10 or conversion rates ≥ 5% in the control sample were removed. We calculated the RNA A/T to G/C editing frequency for all A/T bases in the mitochondrial genome and plotted mitochondrial genome-wide graphs (Supplementary Fig. S10).

### Cell lysis for genomic DNA analysis

After removing the growth medium, HEK 293T cells were treated with 100LμL of cell lysis buffer (50LmM Tris-HCl; pH 8.0, 1LmM EDTA, 0.005% sodium dodecyl sulfate) supplemented with 5LμL of Proteinase K (Qiagen). The cells were lysed by incubation at 55L°C for 1Lh, and then at 95L°C for 10Lmin. The genomic DNA mixture was subjected to targeted deep sequencing.

### Targeted deep sequencing

NGS libraries for targeted deep sequencing were created using nested PCR. The target area was initially amplified by PCR using PrimeSTAR® GXL polymerase (Takara). Amplicons were amplified again by PCR with TruSeq DNA-RNA CD index-containing primers to label each fragment with adapter and index sequences in order to build NGS libraries. The PCR primers are listed in Table S1. The final PCR products were purified with a PCR purification kit (MGmed) and sequenced using a MiniSeq sequencer (Illumina). Base editing frequencies from targeted deep sequencing data were measured using source code (https://github.com/ibs-cge2/prime_editor_analysis, written by BotBot Inc.), as previously described^57^.

### RNA purification from cultured mammalian cells and targeted RNA sequencing

RNA was extracted from cultured cells using a NucleoSpin RNA Plus kit (Macherey-Nagel) according to the manufacturer’s instructions. RNAs were then reverse transcribed to create cDNAs using a SuperScript IV Reverse Transcriptase Kit (Thermo Fisher), again following the manufacturer’s instructions. Regions of interest were then amplified by PCR. Amplified regions were sequenced using the targeted deep sequencing procedure described above.

### Relative ratio of DNA on-target editing frequencies to RNA off-target editing frequencies

To analyze the DNA on-target editing frequency of variants against the RNA off-target editing frequency at six representative sites, the DNA on-target activity and RNA off-target activity of the variants were normalized to the sTALED values. Then, the normalized DNA on-target value was divided by the normalized RNA off-target value, as shown below.

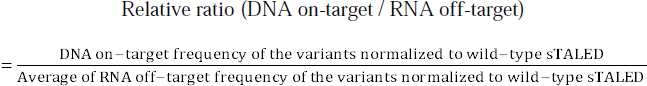

In the case of wild-type sTALED, this value is 1 because both the normalized DNA on-target value and RNA off-target value are 1. A higher relative ratio indicates lower off-target activity compared to the on-target activity.

### Whole mitochondrial genome sequencing

For whole mitochondrial genome sequencing, three procedures were required: PCR amplification, NGS library creation, and NGS. Initially, cells were treated with 100LμL of cell lysis buffer (50LmM Tris-HCl; pH 8.0, 1LmM EDTA, 0.005% sodium dodecyl sulfate) supplemented with 5LμL of Proteinase K (Qiagen) after the growth medium was removed. The cells were lysed by incubation at 55L°C for 1Lh, and then at 95L°C for 10Lmin. mtDNA was then amplified by PCR using PrimeSTAR® GXL polymerase (Takara). PCR was performed using two sets of slightly overlapping primers to reduce primer bias. Each primer pair amplified almost 50% of the mtDNA. The PCR products were then purified with a PCR purification kit (MGmed). Finally, we utilized an Illumina DNA Prep kit with Nextera DNA CD Indexes to create an NGS library from the purified PCR products (Illumina). The libraries were then pooled and transferred onto a MiniSeq sequencer (Illumina).

### Analysis of mitochondrial genome-wide off-target editing

To analyze NGS data from whole mitochondrial genome sequencing, we adopted a method previously used to investigate off-target effects in mitochondrial genomes^18–20^. Initially, we aligned the Fastq sequences to the GRCh38 (release v102) reference genome using BWA (v.0.7.17), and then created BAM files with SAMtools (v.1.9) by fixing read pairing information and flags. Subsequently, we utilized the REDItoolDenovo.py script from REDItools (v.1.2.1) to find all thymines and adenines in the mitochondrial genome with conversion rates > 0.1%. Positions with conversion rates ≥ 10% in both treated and untreated samples were identified as SNVs in the cell lines and removed. We also excluded the construct’s on-target sites. The remaining sites were regarded as off-target sites, and we counted the number of edited A/T nucleotides with an editing frequency > 0.1%. We calculated the average A/T to G/C editing frequency for all bases in the mitochondrial genome by averaging the conversion rates at each base location in the off-target sites, as shown below.

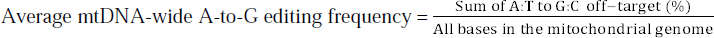

Mitochondrial genome-wide graphs were constructed by plotting the conversion rates at on-target and off-target sites with an editing frequency ≥ 1% across the entire mitochondrial genome.

### Cell viability assays

Cell viability assays were performed using CellTiter 96® Aqueous One Solution (Promega) at day 2 or day 4 after plasmid transfection. Cell viability assay measures the number of viable cells with a colorimetric method. Cells were treated with CellTiter 96® Aqueous One Solution, and the quantification of bio-reduced product was measured by recording the absorbance at 490 nm according to the manufacturer’s instructions.

### Echocardiography analysis

Echocardiography was performed on *mtATP6* mutant mice, with same-aged non-edited mice used as a control. Before echocardiography, body weights were measured on electronic scales (Satorius), and mice were anesthetized using 1.2% avertin with the dosage based on their body weight. Echocardiographic observations were performed at the same time after anesthesia with a Vivid™ iq Ultrasound system (GE HealthCare). Experimental procedures and data analysis were conducted based on the International Mouse Phenotyping Consortium (IMPC) standards (www.mousephenotype.org).

## DATA AVAILABILITY

All data supporting the results are available in the main text or supplementary materials. The data that support the findings of this study are available from the corresponding author upon request. The high-throughput sequencing data from this study have been deposited in the NCBI Sequence Read Archive (SRA) database under the accession code PRJNA947479.

## CODE AVAILABILITY

Base editing frequencies and indel frequencies from targeted deep sequencing data were calculated with source code (https://github.com/ibs-cge2/prime_editor_analysis) ^57^.

## REFERENCES

1. Silva-Pinheiro, P. & Minczuk, M. The potential of mitochondrial genome engineering. Nat Rev Genet (2021).

2. Shoop, W.K., Bacman, S.R., Barrera-Paez, J.D. & Moraes, C.T. Mitochondrial gene editing. Nature Reviews Methods Primers 3, 19 (2023).

3. Falabella, M., Minczuk, M., Hanna, M.G., Viscomi, C. & Pitceathly, R.D.S. Gene therapy for primary mitochondrial diseases: experimental advances and clinical challenges. Nat Rev Neurol 18, 689–698 (2022).

4. Di Donfrancesco, A. et al. Gene Therapy for Mitochondrial Diseases: Current Status and Future Perspective. Pharmaceutics 14 (2022).

5. Zhong, G., Venkatesan, J.K., Madry, H. & Cucchiarini, M. Advances in Human Mitochondria-Based Therapies. Int J Mol Sci 24 (2022).

6. Kar, B., Castillo, S.R., Sabharwal, A., Clark, K.J. & Ekker, S.C. Mitochondrial Base Editing: Recent Advances towards Therapeutic Opportunities. Int J Mol Sci 24 (2023).

7. Phan, H.T.L., Lee, H. & Kim, K. Trends and prospects in mitochondrial genome editing. Exp Mol Med 55, 871–878 (2023).

8. Sabharwal, A. et al. A Primer Genetic Toolkit for Exploring Mitochondrial Biology and Disease Using Zebrafish. Genes (Basel) 13 (2022).

9. Khotina, V.A., Vinokurov, A.Y., Bagheri Ekta, M., Sukhorukov, V.N. & Orekhov, A.N. Creation of Mitochondrial Disease Models Using Mitochondrial DNA Editing. Biomedicines 11 (2023).

10. Lee, S. et al. Enhanced mitochondrial DNA editing in mice using nuclear-exported TALE-linked deaminases and nucleases. Genome Biol 23, 211 (2022).

11. Lee, H. et al. Mitochondrial DNA editing in mice with DddA-TALE fusion deaminases. Nat Commun 12, 1190 (2021).

12. Jiayin Guo, X.C., Zhiwei Liu, Haifeng Sun, Yu Zhou, Yichen Dai, Yu’e Ma, Lei He, Xuezhen Qian, Jianying Wang, Jie Zhang, Yichen Zhu, Jun Zhang, Bin Shen, and Fei Zhou DdCBE mediates efficient and inheritable modifications in mouse mitochondrial genome. Molecular Therapy Nucleic Acids VOLUME 27, P73–80 (2022).

13. Willis, J.C.W., Silva-Pinheiro, P., Widdup, L., Minczuk, M. & Liu, D.R. Compact zinc finger base editors that edit mitochondrial or nuclear DNA in vitro and in vivo. Nat Commun 13, 7204 (2022).

14. Chen, X. et al. DdCBE-mediated mitochondrial base editing in human 3PN embryos. Cell Discov 8, 8 (2022).

15. Tan, L. et al. A conditional knockout rat resource of mitochondrial protein-coding genes via a DdCBE-induced premature stop codon. Sci Adv 9, eadf2695 (2023).

16. Taylor, R.W. & Turnbull, D.M. Mitochondrial DNA mutations in human disease. Nat Rev Genet 6, 389–402 (2005).

17. Gorman, G.S. et al. Prevalence of nuclear and mitochondrial DNA mutations related to adult mitochondrial disease. Ann Neurol 77, 753–759 (2015).

18. Mok, B.Y. et al. A bacterial cytidine deaminase toxin enables CRISPR-free mitochondrial base editing. Nature 583, 631–637 (2020).

19. Cho, S.I. et al. Targeted A-to-G base editing in human mitochondrial DNA with programmable deaminases. Cell 185, 1764–1776 e1712 (2022).

20. Lim, K., Cho, S.I. & Kim, J.S. Nuclear and mitochondrial DNA editing in human cells with zinc finger deaminases. Nat Commun 13, 366 (2022).

21. Mok, Y.G. et al. Base editing in human cells with monomeric DddA-TALE fusion deaminases. Nat Commun 13, 4038 (2022).

22. Lee, S., Lee, H., Baek, G. & Kim, J.S. Precision mitochondrial DNA editing with high-fidelity DddA-derived base editors. Nat Biotechnol 41, 378–386 (2023).

23. Mok, B.Y., et al. CRISPR-free base editors with enhanced activity and expanded targeting scope in mitochondrial and nuclear DNA. Nat Biotechnol 40, 1378–1387 (2022).

24. Mi, L. et al. DddA homolog search and engineering expand sequence compatibility of mitochondrial base editing. Nat Commun 14, 874 (2023).

25. Yin, L., Shi, K. & Aihara, H. Structural basis of sequence-specific cytosine deamination by double-stranded DNA deaminase toxin DddA. Nat Struct Mol Biol (2023).

26. Guo, J., et al. A DddA ortholog-based and transactivator-assisted nuclear and mitochondrial cytosine base editors with expanded target compatibility. Mol Cell 83, 1710–1724 e1717 (2023).

27. Richter, M.F., et al. Phage-assisted evolution of an adenine base editor with improved Cas domain compatibility and activity. Nat Biotechnol 38, 883–891 (2020).

28. Gaudelli, N.M., et al. Directed evolution of adenine base editors with increased activity and therapeutic application. Nat Biotechnol 38, 892–900 (2020).

29. Jeong, Y.K., et al. Adenine base editor engineering reduces editing of bystander cytosines. Nat Biotechnol 39, 1426–1433 (2021).

30. Gaudelli, N.M. et al. Programmable base editing of A*T to G*C in genomic DNA without DNA cleavage. Nature 551, 464–471 (2017).

31. Gammage, P.A., Moraes, C.T. & Minczuk, M. Mitochondrial Genome Engineering: The Revolution May Not Be CRISPR-Ized. Trends Genet 34, 101–110 (2018).

32. Grunewald, J., et al. Transcriptome-wide off-target RNA editing induced by CRISPR-guided DNA base editors. Nature 569, 433–437 (2019).

33. Zhou, C. et al. Off-target RNA mutation induced by DNA base editing and its elimination by mutagenesis. Nature 571, 275–278 (2019).

34. Grunewald, J., et al. CRISPR DNA base editors with reduced RNA off-target and self-editing activities. Nat Biotechnol 37, 1041–1048 (2019).

35. Rees, H.A., Wilson, C., Doman, J.L. & Liu, D.R. Analysis and minimization of cellular RNA editing by DNA adenine base editors. Sci Adv 5, eaax5717 (2019).

36. Li, J., et al. Structure-guided engineering of adenine base editor with minimized RNA off-targeting activity. Nat Commun 12, 2287 (2021).

37. Zhang, Z., et al. Engineering an adenine base editor in human embryonic stem cells with minimal DNA and RNA off-target activities. Mol Ther Nucleic Acids 29, 502–510 (2022).

38. Chen, L., et al. Engineering a precise adenine base editor with minimal bystander editing. Nat Chem Biol 19, 101–110 (2023).

39. Lapinaite, A., et al. DNA capture by a CRISPR-Cas9-guided adenine base editor. Science 369, 566–571 (2020).

40. Moslemi, A.R., Darin, N., Tulinius, M., Oldfors, A. & Holme, E. Two new mutations in the MTATP6 gene associated with Leigh syndrome. Neuropediatrics 36, 314–318 (2005).

41. Chen, B., Daneshgar, N., Lee, H.C., Song, L.S. & Dai, D.F. Mitochondrial Oxidative Stress Mediates Bradyarrhythmia in Leigh Syndrome Mitochondrial Disease Mice. Antioxidants (Basel) 12 (2023).

42. Mok, Y.G., Hong, S., Bae, S.J., Cho, S.I. & Kim, J.S. Targeted A-to-G base editing of chloroplast DNA in plants. Nat Plants 8, 1378–1384 (2022).

43. Li, R. et al. High-efficiency plastome base editing in rice with TAL cytosine deaminase. Mol Plant 14, 1412–1414 (2021).

44. Kang, B.C. et al. Chloroplast and mitochondrial DNA editing in plants. Nat Plants 7, 899–905 (2021).

45. Guo, J. et al. Precision modeling of mitochondrial diseases in zebrafish via DdCBE-mediated mtDNA base editing. Cell Discov 7, 78 (2021).

46. Panja, C., Niedzwiecka, K., Baranowska, E., Poznanski, J. & Kucharczyk, R. Analysis of MT-ATP8 gene variants reported in patients by modeling in silico and in yeast model organism. Sci Rep 13, 9972 (2023).

47. Baranowska, E. et al. Probing the pathogenicity of patient-derived variants of MT-ATP6 in yeast. Dis Model Mech 16 (2023).

48. Zhang, S., et al. TadA reprogramming to generate potent miniature base editors with high precision. Nat Commun 14, 413 (2023).

49. Chen, L. et al. Re-engineering the adenine deaminase TadA-8e for efficient and specific CRISPR-based cytosine base editing. Nat Biotechnol 41, 663–672 (2023).

50. Zheng, S. et al. Efficient PAM-Less Base Editing for Zebrafish Modeling of Human Genetic Disease with zSpRY-ABE8e. J Vis Exp (2023).

51. Xue, N., et al. Improving adenine and dual base editors through introduction of TadA-8e and Rad51DBD. Nat Commun 14, 1224 (2023).

52. Ryu, S.M. et al. Adenine base editing in mouse embryos and an adult mouse model of Duchenne muscular dystrophy. Nat Biotechnol 36, 536–539 (2018).

53. Newby, G.A. et al. Base editing of haematopoietic stem cells rescues sickle cell disease in mice. Nature 595, 295–302 (2021).

54. Suh, S. et al. Restoration of visual function in adult mice with an inherited retinal disease via adenine base editing. Nat Biomed Eng 5, 169–178 (2021).

55. Kim, Y. et al. A library of TAL effector nucleases spanning the human genome. Nat Biotechnol 31, 251–258 (2013).

56. Khanna, A., et al. Bam-readcount -- rapid generation of basepair-resolution sequence metrics. ArXiv (2021).

57. Lee, J. et al. Prime editing with genuine Cas9 nickases minimizes unwanted indels. Nat Commun 14, 1786 (2023).

